# Predicting RNA Sequence-Structure Likelihood via Structure-Aware Deep Learning

**DOI:** 10.1101/2024.01.04.574208

**Authors:** You Zhou, Giulia Pedrielli, Fei Zhang, Teresa Wu

## Abstract

**Motivation:** The active functionalities of RNA are recognized to heavily dependent on the structure and sequence. Therefore, A model that can accurately evaluate a design by giving RNA sequence-structure pairs would be a valuable tool for many researchers. Machine learning methods have been explored to develop such tools, showing promising results. However, two key issues remain. Firstly, the performance of machine learning models is affected by the features used to characterize RNA. Currently, there is no consensus on which features are the most effective for characterizing RNA sequence-structure pairs. Secondly, most existing machine learning methods extract features describing entire RNA molecule. We argue that it is essential to define additional features that characterize nucleotides and specific sections of RNA structure to enhance the overall efficacy of the RNA design process.

**Results:** We develop two deep learning models for evaluating RNA sequence-structure pairs. The first model, NU-ResNet, uses a convolutional neural network architecture that solves the aforementioned problems by explicitly encoding RNA sequence-structure information into a 3D matrix. Building upon NU-ResNet, our second deep learning model, NUMO-ResNet, incorporates additional information derived from the characterizations of RNA, specifically the 2D folding motifs. In this work, we introduce an automated method to extract these motifs based on fundamental secondary structure descriptions. To assess the robustness of our models, we conduct 10-fold cross validation. Furthermore, we evaluate the performance of both models on two independent testing datasets. Our proposed models demonstrate excellent performance across both datasets and surpass the performance of the ENTRNA approach.

**Availability and Implementation:** The corresponding source code and data for this research is available at https://github.com/yzhou617/NU-ResNet_and_NUMO-ResNet.

**Supplementary information:** Supplementary data are available at Bioinformatics online.

## Introduction

RNA molecules play an important role in protein synthesis, gene regulation, and catalysis [22, 9]. It is composed of four types of nucleotides, adenine (A), uracil (U), cytosine (C), and guanine (G) [15]. RNA studies have witnessed an important growth in the areas of materials in nanotechnology applications [15]. For example, RNA aptamers can be utilized as biosensors [25], riboswitches that exist in non-coding parts of messenger RNA (mRNA) can control gene expression [37], and RNA strands can be used to construct nanoscaffolds for therapeutic applications in nanomedicine [18]. In many of these applications, the functionality of the designed RNA molecule is highly influenced by its geometrical structure [22, 27]. In general, the structure can be represented in its primary (sequence of nucleotides), secondary (sequence and pairings between nucleotides), and tertiary form (sequence, pairings, and 3D displacement of nucleotides), with increasing associated computational, and experimental complexity. Although RNA tertiary structure can provide insights into the actual 3D geometry, the scope of this study is specifically directed towards the investigation of RNA secondary structure. In fact, RNA secondary structure exhibits greater stability compared to the tertiary structure folding [34]. Moreover, learning RNA secondary structure can help understand and predict the tertiary structure folding [32, 34]. Therefore, developing a comprehensive understanding of RNA secondary structure, along with its associated patterns, can enhance our understanding of RNA and its functional mechanisms.

In this study, we present a series of deep learning models designed to assess RNA sequence-secondary structure pairs, focusing exclusively on pseudoknot-free structures. To describe the distinctive shapes within the secondary structure, we employ the term *motif*, which will be defined later in Section 2. The underlying concept involves using specific sub-structures to capture the localized patterns formed by subsets of nucleotides.

### Background and Motivation

Several scientific contributions have focused on the analysis of RNA secondary structures with three core research areas: *secondary structure prediction*: given a sequence, we can predict the secondary structure the RNA will fold into [21]; *inverse folding prediction*: given a secondary structure, we can predict the most likely sequence to obtain the target structure [14]; (iii) problems (i)-(ii) have in common the need to evaluate the *quality* of a *sequence-structure* pair [33]: given a structure and a sequence, we can evaluate the likelihood of observing the pair [33]. Recently, it has been shown how methods from (iii) can be used in (i) and, potentially, (ii) in the form of *experts* that evaluate intermediate structures [20]. In the following, we briefly review the models adopted by state-of-the-art approaches in (i) and (ii) to evaluate sequence-structure pairs.

Within the field of *RNA secondary structure prediction*, several contributions have focused on the use of minimum free energy (MFE) as a key metric that explains folding. The underlying assumption is that the structure with the lowest free energy is also the most likely structure the RNA will adopt. It is important to highlight that, generally, free energy cannot be calculated in closed form due to the (a) incomplete understanding of the RNA molecular interactions, and (b) the impractical computational cost of detailed kinetic simulation tools. As a result, several approximate models have been proposed in the literature [23, 38, 3, 2] to estimate the free energy associated with a given secondary structure. An example of methods in this class is RNAfold [21], which uses the approach in [42] to approximate MFE. MXfold2 [29], SPOT-RNA [31], and SPOT-RNA2 [32] utilize deep learning (DL) to predict the RNA secondary structure. Specifically, MXfold2 predicts the RNA secondary structure by maximizing a score which is the sum of a DL model generated folding score and the contribution from thermodynamic parameters. In the DL model of MXfold2, the thermodynamic regularization in the loss function prevents the folding score of the DL model from being significantly different from the free energy calculated by thermodynamic parameters. SPOT-RNA employs Transfer Learning [26] where the input is the outer concatenation of the one hot encoding of the RNA sequence, and the output is an upper triangular matrix whose size scales quadratically with the length of the RNA sequence [36, 31]. The non-zero elements of the upper triangular matrix represent the likelihood for each pair of nucleotides to bind, and they are used to predict the RNA secondary structure [31]. The SPOT-RNA2 adds three features to the RNA sequence as the input: the predicted probability of base pairing obtained from a Linear Partition algorithm [40], the Position Specific Score Matrix (PSSM), and the information of Direct Coupling Analysis (DCA) [41, 19]. The output of SPOT-RNA2 is still an upper triangular matrix whose non-zero elements represent the likelihood of being paired for each pair of nucleotides [32]. ExpertRNA is a Reinforcement Learning algorithm that uses the rollout method [7, 8, 5, 6] to predict RNA secondary structure [20]. The tool uses RNAfold to generate multiple For the challenge that there is no consensus on which features are the most effective for characterizing RNA sequence-secondary structure pairs, the NU-ResNet explicitly encodes the RNA sequence-secondary structure pairs by using an innovative image representation, a 3D intermediate candidate structures [21] that are evaluated using the ENTRNA tool presented in [33].

In *RNA inverse folding* area, NUPACK [39] is among the most commonly used methods. NUPACK formulates the inverse folding as an optimization problem whose objective is to minimize the ensemble defect defined as the average of wrongly paired nucleotides’ counts on the ensemble of unpseudoknotted structures. When designing the RNA sequence, NUPACK decomposes the target structure into sub-structures that are then optimized. RNAiFold employs the MFE as the objective to predict the RNA sequence for a given RNA secondary structure [14, 13]. The RNAiFold includes two approaches, a Constraint Programming (CP) [35] based algorithm and Large Neighborhood Search (LNS) based algorithm. The constraints of the CP ensure the solutions can follow the RNA folding rules, possess desired features of RNA design, fold into the corresponding target RNA secondary structures. The only difference between CP and LNS is regarding the search method. The CP searches the entire space while the LNS fixes parts of the solution space and explores the unfixed parts.

From the review we observed how, independently from the applications, *evaluating the RNA sequence-secondary structure pair* is a core element of the methodologies. Example evaluating metrics include the ones used in the partition method [24] and LinearPartition [40]. The partition-based methods rely on the Boltzmann distribution where the molecule with lower energy has higher probability to exist. However, the consensus regarding which metric should be utilized to evaluate RNA secondary structure has not been made [33]. There has been a notable rise in the utilization of machine learning-based approaches that incorporate features beyond MFE-based methods. This is justified by the recognition that RNA molecules often exhibit stability levels higher than what is predicted by MFE calculations [33]. Additionally, considering the folding kinetics becomes important when dealing with large RNAs [22]. In this direction, ENTRNA is a classifier [33] that utilizes domain knowledge to determine features for encoding the information of RNA sequence-structure pairs and develops the machine learning model (i.e. logistic regression) to evaluate the pair of RNA sequence and secondary structure. ENTRNA output score is defined by the authors as *foldability*, and it can be interpreted as the likelihood that the RNA sequence and RNA secondary structure coexist. While very similar to what we want to achieve, ENTRNA presents two key challenges. The first challenge is that the proposed features may not fully express the sequence and structure properties. This is because there is no agreement on which features should be used to characterize the RNA folding. In addition, the extracted features are in RNA level, which may not thoroughly encode rich information of nucleotides and sub-structures.

### Contribution and Paper Structure

In order to tackle the aforementioned challenges and create a new metric to evaluate the co-existence of a RNA sequence-structure pair, we propose a deep learning based approach including two deep learning models, NU-ResNet and NUMO-ResNet. Our contributions are:

1. For the challenge that there is no consensus on which features are the most effective for characterizing RNA sequence-secondary structure pairs, the NU-ResNet explicitly encodes the RNA sequence-secondary structure pairs by using an innovative image representation, a 3D matrix. Thus, a convolutional neural networks (CNN), ResNet-18 [16], can be employed to automatically extract features from our proposed 3D matrix.
2. In the face of the challenge that the features extracted in existing machine learning method characterize the entire RNA molecule (e.g. GC percentage). Our proposed 3D matrix is designed to incorporate the nucleotide level information including nucleotides’ types and their base pairing. The NUMO-ResNet extends the NU-ResNet by incorporating sub-structure (i.e. motif) information including motifs’ types and their free energy, which is encoded into a 2D matrix. We revise the architecture based on ResNet-18 to take both of our proposed 3D matrix and 2D matrix as inputs to develop NUMO-ResNet.

To our knowledge, this is the first paper which utilizes neural networks to evaluate the pair of RNA sequence and RNA secondary structure.

The performance of NU-ResNet and NUMO-ResNet outperform state-of-the-art model, ENTRNA, in two testing data sets which are totally untouched during the training. In addition, the extended model, NUMO-ResNet, significantly improves the prediction accuracy based on 10-fold cross validation (CV) results and the convergence behavior during the training compared to the NU-ResNet.

The remainder of the paper is structured as follows: in Materials and Methods section, we elaborate the detailed algorithms for encoding the pair of RNA sequence and RNA secondary structure as well as the two neural network architectures utilized in our proposed framework. The section Results introduces the metrics used to investigate the models’ performance, the data sets utilized in this research, the models’ performance comparison, and models’ convergence behavior during the training process. In section Conclusion, we summarize the work and introduce the potential directions for future research.

## Materials and Methods

Fig. 1 shows an example of RNA secondary structure represented as a graph using the software VARNA [11]. The RNA graph *G*_RNA_ = (*V, E, F*) has vertices *V*, edges *E*, and faces *F*. Each element in the set of vertices *v* ∈ *V* has an associated label *𝓁* (*v*) ∈ {*C, G, A, U* }, based on its composition being, cytosine, guanine, adenine, and uracil, respectively. Edges *e* ∈ *E* of *G*_RNA_ are of two types: *hydrogen bonds* connecting two paired nucleotides (we refer to this subset of edges as *E*_*H*_), also commonly referred to as *interior edges*; and *phosphodiester bonds* connecting any two adjacent nucleotides (we refer to this subset of edges as *E*_*P*_), commonly referred to as *exterior edges*. As a result, *E* = *E*_*H*_ ∪ *E*_*P*_. All edges in *E* are labelled using the dot-bracket notation presented in [4]. An interior edge is represented by a pair of open and closed brackets (“(“,”)”), and an exterior edge is represented by two consecutive dots (“.”). The details of the dot-bracket notation of pseudoknot-free RNA are introduced in the supplementary file.

**Fig. 1.**
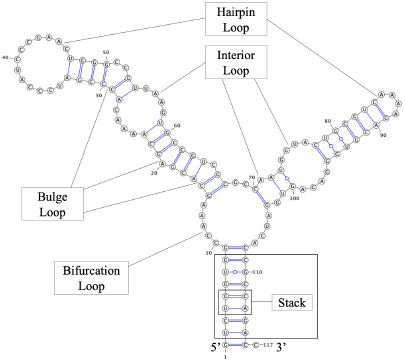
An example of RNA represented by RNA secondary structure. This RNA is 5s Paracoccus-denitrificans-1, extracted from the Mathews laboratory data set [28]. The 5 motifs considered in our approach are indicated: hairpin, interior, bulge, bifurcation loops, and stack.

Finally, each element *f* ∈ *F*, a face, is the 2-d region defined by the tuple 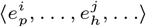, i.e., a closed region bounded by consecutive hydrogen and phosphodiester bonds. The number and arrangement of edges determines the size and the shape of each face. Given the constraints that nucleotides have to satisfy when bonding, 5 shapes exist for faces in 2-d RNA structures (see Fig. 1 for a depiction). In the rest of the paper, we refer to the shape of the faces as *motifs*. Please see the definition of these 5 motifs in the supplementary file.

The corresponding RNA sequence and RNA secondary structure in dot-bracket notation of graph form by Fig. 1 are introduced in the supplementary file.

Section 2.1 introduces our first Deep Learning (DL) model in which the nucleotide level information of an RNA sequence-structure pair is encoded into a 3D matrix. We name this model NU-ResNet. Section 2.2 introduces our second DL model, which adds motifs to the features encoded by NU-ResNet, we refer to this as NUMO-ResNet.

### Nucleotide-level features-informed Residual Neural Network Model (NU-ResNet)

In this section, we introduce our first deep learning model, which uses nucleotide level properties to evaluate sequence-structure pairs. We refer to this model as NU-ResNet since it uses a Residual Neural Network as basis architecture and it uses nucleotide level features to encode the input.

#### Input Layer Encoding

A key contribution of this work is the design and implementation of the input layer encoding, which we detail herein. The input layer for the first model describes the structure of the molecule. We encode the secondary structure as a 3D matrix of size [*L* × *L* × *B*], where *L* is the length of RNA sequence (number of nucleotides), *B* = 4 is the number of bits we need for the one-hot encoding of the nucleotide type (C,G,A,U), and the base pairing information, we will refer to these as the channels for the input of our NU-ResNet. As a result, a cell in the 3D matrix with index (*i, j*), *i* = 1, …, *L j* = 1, …, *L* is a 4-elements vector with each element in {0, 1}. A key motivation behind the choice of transforming the one-hot encoding into a 3D matrix was driven by the consideration that deep learning approaches are particularly effective with imaging data which are essentially 2D or 3D matrixes, in that they are designed to extract features from this type of input format. In the following, we detail the construction of the 3D matrix starting from input sequence-structure information. The 3D matrix **G** is initialized with all zeros.

#### Diagonal Elements Encoding

We set

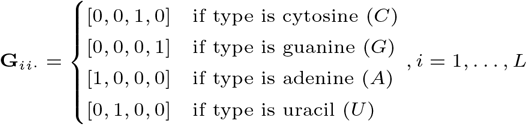

#### Out-of-diagonal Elements Encoding

Next, we encode all the edges *e* ∈ *E*_*H*_ as vectors of the 3D matrix, denoted as **G**_*ij·*_, which satisfy (*i, j*) ∈ *E*_*H*_, i.e., all the nucleotides that form a base pair.

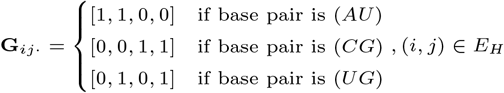

Specifically, the values are obtained as the sum of the one-hot encoding vectors of the two nucleotides involved in the pairing. For symmetry, (*i, j*) ∈ *E*_*H*_ implies (*j, i*) ∈ *E*_*H*_, hence the vector **G**_*ij·*_ is equivalent to the vector **G**_*ji·*_, ∀*i, j* = 1, …, *L*, where the equality between vectors is interpreted as the equality of all elements.

**Example**. In order to clarify the approach, we show an example that leads to the generation of the 3D RNA matrix. We start considering a fictitious RNA sequence “CAGGAGCUCUUC” with corresponding secondary structure “.(((…)))..”, i.e., it contains 3 base pairs.

The visualization of the 3D RNA matrix from this example is shown in Fig. 2.

**Fig. 2.**
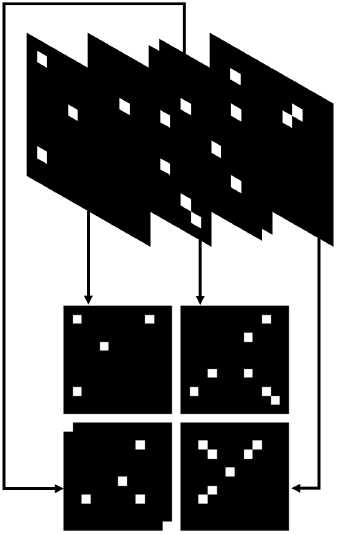
Example of 3D RNA matrix. The combination of four stacked subfigures on the top is the input of NU-ResNet. The flat form of four stacked subfigures are shown at the bottom. The 4 2D matrixes at the bottom represent the encoded *G*_*ij*1_, *G*_*ij*2_, *G*_*ij*3_, and *G*_*ij*4_, respectively. In this 3D RNA matrix example, the white box and black box correspond to 1 and 0 values of the matrix, respectively.

Deep Learning Architecture Design and Architecture Training. Convolutional Neural Networks (CNN) have shown very good performance within the image recognition literature [1, 30, 16]. Within this broad family of learning models, ResNets are designed to resolve the degradation of training accuracy when the depth of the neural network increases [16]. ResNets have shown robust performance on image classification tasks. Rather than learning the sophisticated functions to depict the relationship between input and output directly, ResNets learn how to approximate residual functions by using stacked nonlinear layers [16]. For these reasons, we choose the ResNet-18 as the primary architecture in this research. The Fig. 3 shows the architecture of NU-ResNet. The details of NU-ResNet architecture are introduced in the supplementary file.

**Fig. 3.**
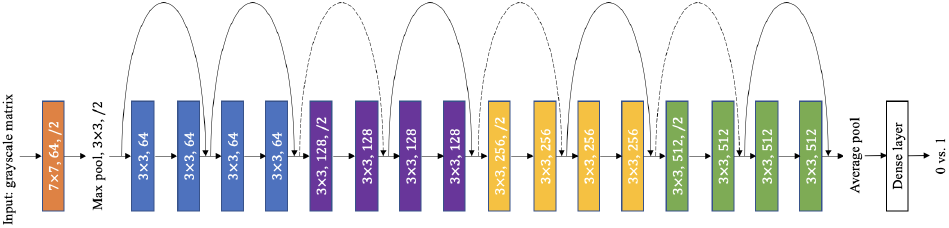
The architecture of NU-ResNet.

The model is trained using RNA molecules of different lengths each corresponding to 3D RNA matrixes of different sizes (*L*). To allow training, we transform the input by “padding” with zeros along each of the *B* = 4 channels. After padding, the size of each 3D RNA matrix equals to ℒ × ℒ× 4, where ℒ is any value greater than the size of the longest RNA we are building the model to evaluate. This value can be given as input by the user or be set to the size of longest sequence across the training, testing, and validation datasets.

### Nucleotide-level features and Motifs-informed Residual Neural Network Model (NUMO-ResNet)

As mentioned in section 1.1, several approaches account for RNA features that knowingly impact the folding. NUMO-ResNet extends the fully data driven (black-box) model in section 2.1 to include features *localized* to sub-structures of the molecule that can potentially impact the RNA stability. In particular, we include the motifs present in the structure, and characterize them with the associated Minimum Free Energy (MFE), and the motif types (see Section 1.1 for the definition of MFE and Section 2, Fig. 1 for the definition of the motifs).

#### Input Layer Encoding

In the following, we detail the approach to automatically derive the motifs contained in the structure, and how to calculate the free energy associated to the different sub-structures given the type of motif and the number of nucleotides involved. The nucleotide type is encoded in the same way as in NU-ResNet.

#### Automatic Identification and Encoding of the Motifs

NUMO-ResNet adds to the base types used in NU-ResNet, the motifs associated to each nucleotide (e.g., stack, hairpin loop, interior loop, bulge loop, bifurcation loop, and no motif). Since a motif involves a face (sub-structure) of a molecule, based on its location within the sub-structure, the same nucleotide may be involved in two motifs. For this reason, for each nucleotide *i* = 1, …, *L*, we can assign two distinct vectors to encode all the possible motifs the base is involved in. This results into two matrixes 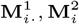 encoded as follows:

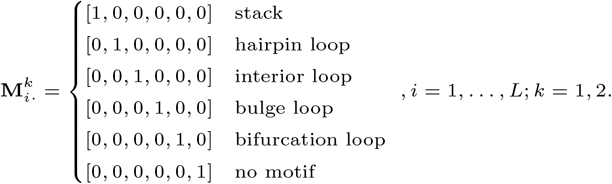

While the encoding of the nucloetide type is a direct translation from the input sequence information, an algorithm is necessary to recover the type and characteristics of the motifs given the sequence and structure information. We develop a procedure to automatically extract motif information for each nucleotide in RNA sequence. Please see the details of this procedure in the supplementary file.

#### Calculating and Encoding Free Energy

The free energy of a molecule is the sum of the free energies calculated at the sub-structure (motif) level [42]. In particular, the free energy of a motif in RNA is a function of the involved bases. In our model, the type and the free energy of a motif are its two core features. Because the motif is composed of nucleotides, we assign the type and the free energy of a motif as the features for each nucleotide that forms it. Since a nucleotide could belong to two adjacent motifs, both of these two adjacent motifs’ types and free energy values should be assigned to this nucleotide as features. To summarize all these features, we propose a 2D matrix, which we refer to as the *nucleotide localized information matrix*, to encode the motifs’ free energy, motifs’ types, and nucleotides’ types for each nucleotide in the RNA molecule. The size of nucleotide localized information matrix is *L* × 5, where *L* is the length of RNA sequence and 5 is the number of motif-driven features.

After representing the categorical variables by one hot encoding, the nucleotide localized information matrix becomes a *L* × 18 matrix (Please see the details in the supplementary file). We pad 0 in the bottom of each nucleotide localized information matrix to uniform the size of all nucleotide localized information matrixes. As a result, each nucleotide localized information matrix has a size of ℒ× 18 (ℒ has the same meaning as defined in section 2.1). In the supplementary file, we introduce how we rescale the numerical variable, free energy.

**Example**. An example of nucleotide localized information matrix is shown in Table 1. The fictitious RNA used in this example is same with the one used in Fig. 2.

**Table 1.** An example of nucleotide localized information matrix. In this table, “nt” is the abbreviation of the nucleotide.

| nt | motif1 | motif1 FE | motif 2 | motif2 FE |
| --- | --- | --- | --- | --- |
| C | none | 0 | none | 0 |
| A | stack | -210 | none | 0 |
| G | stack | -150 | stack | -210 |
| G | hairpin loop | 590 | stack | -150 |
| A | hairpin loop | 590 | none | 0 |
| G | hairpin loop | 590 | none | 0 |
| C | hairpin loop | 590 | none | 0 |
| U | hairpin loop | 590 | stack | -150 |
| C | stack | -150 | stack | -210 |
| U | stack | -210 | none | 0 |
| U | none | 0 | none | 0 |
| C | none | 0 | none | 0 |

#### Architecture Design

To account for the novel features, we revise the architecture proposed for NU-ResNet. Specifically, we have two inputs, 3D RNA matrix and nucleotide localized information matrix, for each RNA. We develop a neural network model which uses two parallel ResNet-18 with removing the last fully connected layer to generate features from each input and concatenates these generated features. The concatenated features are followed by stacked fully connected layers to perform classification. We name this revised ResNet-18 as NUMO-ResNet. The detailed architecture of NUMO-ResNet is shown in Fig. 4.

**Fig. 4.**
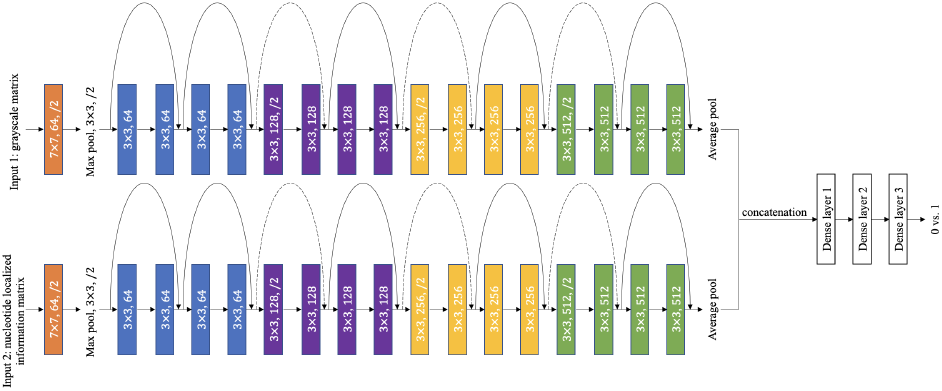
The details of architecture of NUMO-ResNet.

## Results

In this section, we analyze the performance of NU-ResNet and NUMO-ResNet, and analyze them against the state-of-the-art model ENTRNA presented in [33]. Section 3.1 introduces the metrics that are utilized to assess the performance of the different models. Section 3.2 characterizes the datasets utilized in this research, while section 3.3 presents the comparison of performance between NU-ResNet, NUMO-ResNet, and ENTRNA.

### Performance Metrics

Since NU-ResNet and NUMO-ResNet are trained as binary classification models, we propose several metrics to comprehensively analyze the trained architectures. The metrics we utilize include the area under curve receiver operator characteristic (AUCROC), the matthews correlation coefficient (MCC), accuracy, precision, recall, and specificity. The AUCROC has a threshold invariant characteristic which can comprehensively evaluate the classification model. In addition, AUCROC has been proven to be equivalent to the probability that a randomly chosen positive sample can be ranked higher than a randomly chosen negative sample by the classification model [12]. Here, 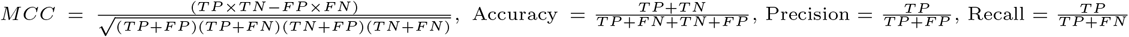,, and 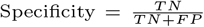. Within these formulas, *TP, TN, FP*, and *FN* refer to the number of true positive, true negative, false positive, and false negative respectively. AUCROC, accuracy, precision, recall, and specificity, are all defined in the range [0, 1]. The MCC metric is defined in the range [−1, 1]. For all of the metrics, higher value indicates better performance.

### Data Sets

The samples utilized in this research are extracted from the data set established by the Mathews laboratory [28], and the Rfam data set [10]. We choose to utilize the samples from these two data sets in this research because they are commonly used in the literature. The longest RNA sequence within the Mathews data set has 338 nucleotides, while the longest sample within Rfam has 408 nucleotides. To unify the size of inputs of NU-ResNet and NUMO-ResNet, we pad both the 3D RNA matrix and the nucleotide localized information matrix with zero elements. We choose to use 410 as the maximum length L, resulting in 3D RNA matrixes with size [410 × 410 × 4] and nucleotide localized information matrixes with size [410 × 18]. When utilizing the CDF of normal distribution to rescale free energy to the [0, 1] interval in the nucleotide localized information matrix, we set *μ* = 0 and *σ* = 5.

Both our deep learning models require positive and negative samples to be trained. The positive samples are the RNA sequence-structure pairs in the data sets, while we generate the negative samples for the same RNA sequences, and we use RNAfold to produce a structure [21] that differs from the true RNA secondary structure in the data set. As a result, each RNA sequence has associated both a *positive sample* and a *negative sample* structure if the RNAfold prediction differs from the true structure. Otherwise, the RNA input only has a positive sample when its RNAfold predicted RNA secondary structure is same with its correct RNA secondary structure. In this work, the Mathews sequence structure pairs are randomly selected to generate the training (TrDS, with 70% of the inputs), validation (VDS, with 15% of the inputs), and testing (TeDS, with 15% of the inputs) datasets. Rfam is used as an extra source for blind testing. Specifically, we eliminate the RNA sequences in Rfam with a CD-HIT-EST-2D similarity score above 0.8 [17].

In the following, we analyze the performance of both our DL models with the associated largest validation accuracy, and lowest validation loss. For these models, we also show the 10-fold CV performance under the combined TrDS and VDS datasets.

### Models Comparison

As previously mentioned, we record the models with best validation accuracy and best validation loss resulting from training. The set of parameters of the model with best validation accuracy and best validation loss is referred to as 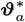and 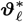respectively. The performance of the resulting NU-ResNet and NUMO-ResNet on the TeDS and Rfam data sets are shown in Table 2 and Table 3. It can be observed how all of four models outperform ENTRNA on both testing data sets.

**Table 2.** Models performance on TeDS. Models with parameters that optimize the validation accuracy are referred to as 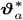, while models with parameters that optimize the validation loss are referred to as 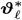.

| Metric | NU-RN |  | NUMO-RN |  | ENTRNA |
| --- | --- | --- | --- | --- | --- |
| | $\vartheta_a^*$ | $\vartheta_\ell^*$ | $\vartheta_a^*$ | $\vartheta_\ell^*$ | |
| Accuracy | 93.12% | 94.94% | 97.17% | 96.76% | 81.78% |
| AUCROC | 0.9736 | 0.9796 | 0.9892 | 0.9885 | 0.8972 |
| MCC | 0.8631 | 0.9002 | 0.9431 | 0.9352 | 0.6560 |
| Precision | 91.01% | 92.47% | 96.97% | 95.90% | 91.75% |
| Recall | 96.56% | 98.47% | 97.71% | 98.09% | 72.14% |
| Specificity | 89.22% | 90.95% | 96.55% | 95.26% | 92.67% |

**Table 3.** Models’ performance on Rfam data set. Models with parameters that optimize the validation accuracy are referred as 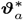, while models with parameters that optimize the validation loss are referred to as 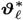.

| Metric | NU-RN |  | NUMO-RN |  | ENTRNA |
| --- | --- | --- | --- | --- | --- |
| | $\vartheta_a^*$ | $\vartheta_\ell^*$ | $\vartheta_a^*$ | $\vartheta_\ell^*$ | |
| Accuracy | 97.60% | 97.31% | 94.91% | 94.91% | 90.72% |
| AUCROC | 0.9734 | 0.9701 | 0.9765 | 0.9800 | 0.9853 |
| MCC | 0.9532 | 0.9475 | 0.9009 | 0.9009 | 0.8209 |
| Precision | 100% | 100% | 98.70% | 98.70% | 86.17% |
| Recall | 95.21% | 94.61% | 91.02% | 91.02% | 97.01% |
| Specificity | 100% | 100% | 98.80% | 98.80% | 84.43% |

In Table 2, we observe that the two NUMO-ResNet models with 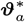or 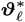outperform ENTRNA on all metrics based on TeDS. In addition, on TeDS, the NU-ResNet with 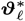 outperforms ENTRNA on 5 out of 6 metrics (i.e. accuracy, AUCROC, MCC, precision, and recall), and the NU-ResNet with 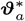outperforms ENTRNA on 4 out of 6 metrics (i.e. accuracy, AUCROC, MCC, and recall).

Table 3 shows that NU-ResNet and NUMO-ResNet outperform ENTRNA on 4 out of 6 metrics (i.e. accuracy, MCC, precision, and specificity) based on the Rfam data set. For AUCROC, the ENTRNA outperforms NU-ResNet and NUMO-ResNet but the difference is in the order of 2% points (i.e. ENTRNA vs. NU-ResNet with 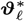). For recall, ENTRNA exceeds our trained models that have lowest recall (i.e. NUMO-ResNet with 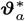 and 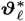) 5.99%. Two saved NUMO-ResNet models outperform ENTRNA on accuracy (4.19%), precision (12.53%), specificity (14.37%), and MCC (0.08).

In Table 2 and 3, ENTRNA outperforms NU-ResNet or NUMO-ResNet in 11 comparisons out of total 48 comparisons, but NU-ResNet and NUMO-ResNet outperform ENTRNA in all remaining 37 comparisons. In most of these 37 comparisons, NU-ResNet and NUMO-ResNet outperform ENTRNA by a large margin. The comparisons where ENTRNA outperforms NU-ResNet or NUMO-ResNet are viewed as the work we are going to work on in the future. Therefore, we conclude that the overall performance of our proposed methods is mostly superior to ENTRNA on both of TeDS and Rfam data set. In addition, the experiments indicate the effectiveness of the methods utilized by NU-ResNet and NUMO-ResNet to encode the RNA sequence-structure pair.

When training NU-ResNet and NUMO-ResNet, we employ Adam as the optimizer of the models parameters. The learning rate and weight decay of optimizer for NU-ResNet are 0.0001 and 0.15, respectively, and the learning rate and weight decay of optimizer for NUMO-ResNet are 0.0001 and 0.1, respectively. For both of NU-ResNet and NUMO-ResNet, the learning rate decreases exponentially with gamma value equaling to 0.95, the batch size during the training is 20, and both models are trained for 100 epochs. The models with associated the best validation accuracy and the best validation loss are saved as models to be deployed.

### Comparison Between Proposed Models

Table 2 shows that NUMO-ResNet with either 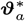or 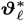 outperforms NU-ResNet with 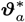 in all 6 metrics. And two saved NUMO-ResNet models outperform NU-ResNet with 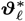 in 5 out of 6 metrics (i.e. accuracy, AUCROC, MCC, precision, and specificity). For recall, the NU-ResNet with 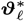 outperforms the NUMO-ResNet with lower recall 0.76%.

Table 3 shows that the two NUMO-ResNet models saved from training have the same performance on almost all metrics except AUCROC. For AUCROC, the NUMO-ResNet with either 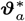 or 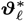 outperforms NU-ResNet models. For the remaining 5 metrics, the NU-ResNet with either 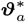 or 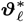outperforms both NUMO-ResNet models. Because of the meaning of AUCROC that is introduced in section 3.1, NUMO-ResNet with 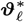 has 98% probability that a randomly chosen positive sample is ranked higher than a randomly chosen negative sample, which indicates that the performance of NUMO-ResNet with 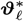 is very good on Rfam data set.

The results of experiments satisfy our expectation because NUMO-ResNet incorporates more features compared to NU-ResNet. Intuitively, NUMO-ResNet should at least has the same performance with NU-ResNet because NUMO-ResNet has the whole input that NU-ResNet has. The results of experiments also show the advance of motif-based features extracted by NUMO-ResNet compared to the input only incorporating sequence and structure information employed by NU-ResNet.

Convergence Behavior of NU-ResNet Training compared to NUMO-ResNet Training.

Here, we provide insights into the training process of the proposed models. In particular, we analyze the validation loss and accuracy metrics as a function of the training effort (i.e., number of epochs). Fig. 5a shows that NUMO-ResNet converges faster compared to NU-ResNet when the training loss is considered. At the same time, NU-ResNet validation loss shows larger fluctuation than NUMO-ResNet. Fig. 5b confirms these observations when accuracy is considered: NUMO-ResNet converges faster in training accuracy compared to NU-ResNet and its validation accuracy is again more stable. These findings suggest that NUMO-ResNet can be trained more efficiently compared to NU-ResNet showing better results also for lower budgets (epochs).

**Fig. 5.**
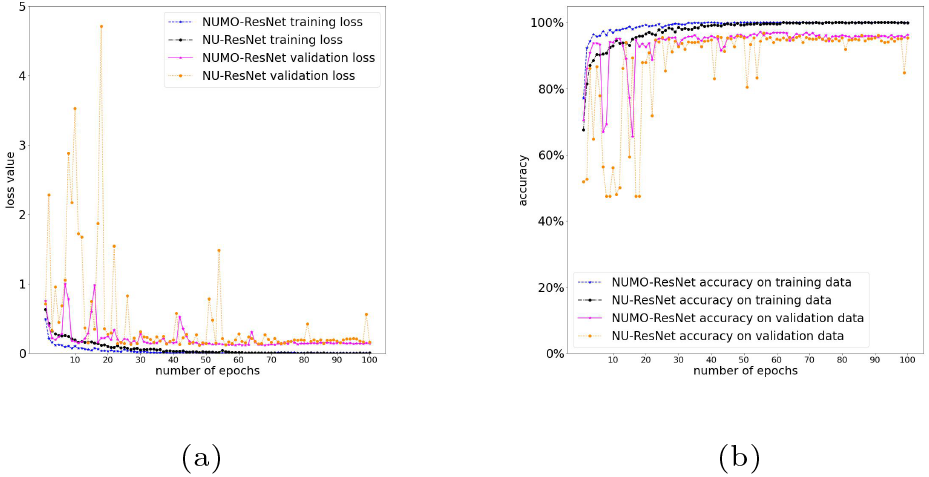
(a): The training and validation loss. (b): The training and validation accuracy.

### Models Robustness Analysis

Since we utilize a weighted sampler to sample the data during the training which has randomness, the performance of trained models on testing data may be affected by this randomness. To verify the robustness of the trained models, we perform a 10-fold CV on both NU-ResNet and NUMO-ResNet. At each iteration of the validation, we use 8 folds of data for training, 1 fold of data for validation, and 1 fold of data for testing. Similar to the previous analysis, for each iteration of the validation routine, we consider two models, one with the best validation accuracy, and one with the best validation loss. Table 4 shows the 10-fold CV results from our models as the average of the performance resulting from 10 iterations of the approach. The 10-fold CV results presented in Table 4 show the consistent performance of the NU-ResNet and NUMO-ResNet across Mathews (Table 2) and Rfam data sets (Table 3).

**Table 4.** 10-fold cross validation results from NU-ResNet and NUMO-ResNet. Models with parameters that optimize the validation accuracy are referred to as 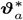, while models with parameters that optimize validation loss are referred to as 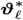.

| Metric | NU-RN |  | NUMO-RN |  |
| --- | --- | --- | --- | --- |
| | $\vartheta_a^*$ | $\vartheta_\ell^*$ | $\vartheta_a^*$ | $\vartheta_\ell^*$ |
| Accuracy | 94.24% | 94.10% | 96.33% | 96.69% |
| AUCROC | 0.9762 | 0.9779 | 0.9880 | 0.9908 |
| MCC | 0.8852 | 0.8819 | 0.9266 | 0.9337 |
| Precision | 93.15% | 93.31% | 96.70% | 96.72% |
| Recall | 96.35% | 95.81% | 96.42% | 97.09% |
| Specificity | 91.84% | 92.16% | 96.22% | 96.23% |

## Conclusion

In this research, we present a deep learning framework for the evaluation of RNA sequence-structure pairs. Within this framework, we introduce two models, NU-ResNet and NUMO-ResNet. Both models exhibit superior performance compared to state-of-the-art RNA sequence-structure pair evaluation models across multiple metrics. The two models rely on different inputs. Particularly, NUMO-ResNet incorporates motif-based features, which enhance the training process efficiency, stability, and overall model performance to a considerable degree. It is important to highlight that this study exclusively focuses on pseudoknot-free structures. Our future efforts will involve addressing pseudoknotted RNA structures, and the outcomes will be reported separately. Furthermore, we plan to explore incorporating uncertainty quantification techniques into our models in future, further enhancing their reliability and robustness.

## Acknowledgments

The authors thank Arizona State University Research Computing for providing us with the computational nodes.

