## Supplemental Material for "Predicting RNA Sequence-Structure Likelihood via Structure-Aware Deep Learning"

### Supplementary Material

#### Introduction of Three Elements in Dot-Bracket Notation

Specifically, the dot-bracket notation of pseudoknot-free RNA is comprised of three elements, namely:

- *dot* (“.”) is used when representing an unpaired nucleotide;
- *open bracket* (“(”) is used when a nucleotide is the origin of a base pair. This nucleotide will be paired with one and only one nucleotide;
- *closed bracket* (“)”) is used when a nucleotide closes a base pair. For symmetry, this nucleotide will be paired with one and only one “open bracket”nucleotide.

As shown below, the information given in graph form by Fig. 1 in the paper can be encoded, without any information loss, using two strings one containing the characters of the bases in the RNA molecule, and the second reporting the dot-bracket notation to express the secondary structure information.

Sequence:
GUCUGGUGGCCAAAGCACGAGCAAAACACCCGAUCCCAUCCCGAACUCGGCCGUUAA
GUGCCGUCGCGCCAAUGGUACUGCGUCAAAAGACGUGGGAGAGUGGAUCACCGCCAG
ACC
Secondary structure:
(((((((((.....(((((.(((....((((((.............))))..))....)))))).))..(.((....(((((((....)))))))....)).)....))))))))).

#### Definition of Motifs

- Stack: A face having two interior edges which are separated by one exterior edge on each side.
- Hairpin loop: A face only having one interior edge.
- Interior loop: A face having two interior edges separated by more than one exterior edge on each side.
- Bulge loop: A face having two interior edges separated by more than one exterior edge on one side and exactly one exterior edge on the other side.
- Bifurcation loop: A face having more than two interior edges.

#### NU-ResNet Architecture

Fig. 1 shows the NU-ResNet architecture with detailing the building block including all the components from the beginning of an identity shortcut to its end, where an identity shortcut is represented by a black solid or dashed curve. The difference between identity shortcuts with black solid and dashed curve is also illustrated in Fig. 1.

#### Procedure to Automatically Extract Motif Information

In the following, we show the key steps of the procedure to automatically extract motif information for each nucleotide in RNA sequence.

- **Initialization**: As previously mentioned, the RNA structure is given as input encoded as a string of dots and brackets, which we refer to as $\Sigma$ of size $L$ (number of nucleotides). We create a set for each motif type and initialize it to the empty set, namely the set of stacks $S^{\text{stack}} = \emptyset$, the set of hairpin loops $S^{\text{hairpin}} = \emptyset$, the set of bulge loops $S^{\text{bulge}} = \emptyset$, the set of interior loops $S^{\text{interior}} = \emptyset$, and the set of bifurcation loops $S^{\text{bifurcation}} = \emptyset$.
- **Step 1.a Base Pair Identification**: All base pairs within $\Sigma$ are translated into a tuple $(j, k), j, k \in \{1, \ldots, L\}$ representing the index of the origin and destination nucleotide, respectively. The collection of all the tuples forms the base pairs index set $\mathbf{B}$ whose elements $\mathbf{b}_i, i = 1, \ldots, |\mathbf{B}|$ are defined as $\mathbf{b}_i = (j, k)$. We adopted the `rna-tools` [3] Python package for the automatic identification of all base pairs within this RNA secondary structure. The implemented algorithm runs with complexity $O(L)$.

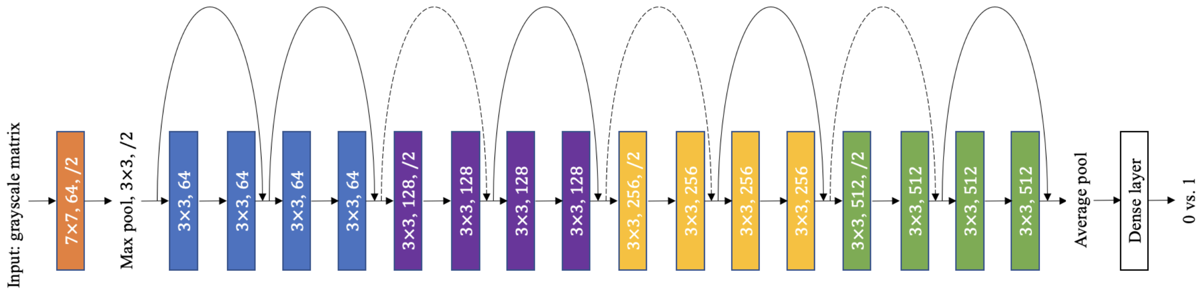


Figure 1: NU-ResNet architecture. The black straight and solid arrows and the curved dashed arrows represent identity shortcuts, additive operators that combines the input and the output of a layer returning the residuals used to learn the model. The black solid curved arrows are used in cases where the input and output have the same dimensionality, and the black dashed curve is used when output dimension is larger [1]. Each identity shortcut corresponds to a *building block* which is a network sub-structure including 2 convolutional layers, 2 Batch Normalization, 2 rectified linear activation units (ReLU) [4] (enabling non-linear models), and 1 identity shortcut. The Batch Normalization [2] is used between each convolutional layer and the ReLU activation to normalize the intermediate representation [2, 1]. The kernel size of the first convolutional layer is 7 and the kernel size of any convolutional layer within the building block is 3. The number of output channels from the convolutional layers in the 4 types of building block is $64, 128, 256$, and $512$, respectively. The parameters in the convolutional layer at the beginning of architecture, in the eight building blocks, and in the fully connected layer (i.e. Dense layer) need to be learned during training.

- **Step 1.b Motif Elicitation:** As previously explained, each motif is bounded by at least one base pair. Hence, given the collection of base pairs from Step 1.a, $\mathbf{B}$, we verify which motifs the base pairs are defined in. More specifically:
  - Stack: If $(\mathbf{b}_i(1)+1, \mathbf{b}_i(2)-1) \in \mathbf{B}$, then

$$S^{\text{stack}} \leftarrow S^{\text{stack}} \cup \{\mathbf{b}_i, (\mathbf{b}_i(1)+1, \mathbf{b}_i(2)-1), \\ (\mathbf{b}_i(1), \mathbf{b}_i(1)+1), (\mathbf{b}_i(2)-1, \mathbf{b}_i(2))\},$$

    where $\{\mathbf{b}_i, (\mathbf{b}_i(1)+1, \mathbf{b}_i(2)-1)\}$ are interior edges (hydrogen bonds), while $\{(\mathbf{b}_i(1), \mathbf{b}_i(1)+1), (\mathbf{b}_i(2)-1, \mathbf{b}_i(2))\}$ are exterior edges (phosphodiester bonds);
  - Hairpin loop: If the structure elements $\Sigma_j =$ ".", $\forall j = \mathbf{b}_i(1)+1, \ldots, \mathbf{b}_i(2)-1$, then

$$S^{\text{hairpin}} \leftarrow S^{\text{hairpin}} \cup \{\mathbf{b}_i, (\mathbf{b}_i(1)+1, \mathbf{b}_i(1)+2), \\ \ldots, (\mathbf{b}_i(2)-2, \mathbf{b}_i(2)-1)\},$$

    where all edges are phosphodiester bonds, except the hydrogen bond $\mathbf{b}_i$;
  - Bulge loop: Similar to the stack, we have two consecutive base pairs whose bases are either at the origin or destination. In particular, if the bulge is *on the side of the destination* bases, we have that $\mathbf{b}_i(1)+1 = \mathbf{b}_{i+1}(1)$, with a bulge of size $\mathbf{b}_i(2) - \mathbf{b}_{i+1}(2) > 1$. Then,

$$S^{\text{bulge}} \leftarrow S^{\text{bulge}} \cup \{\mathbf{b}_i, \mathbf{b}_{i+1}, (\mathbf{b}_i(1), \mathbf{b}_{i+1}(1)), \\ (\mathbf{b}_{i+1}(2), \mathbf{b}_{i+1}(2)+1), \ldots, (\mathbf{b}_i(2)-1, \mathbf{b}_i(2))\},$$

    where all edges are phosphodiester bonds, except the hydrogen bonds $\mathbf{b}_i, \mathbf{b}_{i+1}$. In case the bulge is *on the side of the origin* bases, we have that $\mathbf{b}_i(2)-1 = \mathbf{b}_{i+1}(2)$, with a bulge of size $\mathbf{b}_{i+1}(1) - \mathbf{b}_i(1) > 1$. Then,

$$S^{\text{bulge}} \leftarrow S^{\text{bulge}} \cup \{\mathbf{b}_i, \mathbf{b}_{i+1}, (\mathbf{b}_i(1), \mathbf{b}_i(1)+1), \\ \ldots, (\mathbf{b}_{i+1}(1)-1, \mathbf{b}_{i+1}(1)), (\mathbf{b}_{i+1}(2), \mathbf{b}_i(2))\},$$

where all edges are phosphodiester bonds, except the hydrogen bonds $\mathbf{b}_i, \mathbf{b}_{i+1}$;

- Interior loop: Similar to the bulge loop, we have two consecutive base pairs whose bases are either at the origin or destination. *On the side of the origin and destination* bases, we have that $\mathbf{b}_{i+1}(1) - \mathbf{b}_i(1) > 1$ and $\mathbf{b}_i(2) - \mathbf{b}_{i+1}(2) > 1$. Then,

$$
\begin{aligned}
S^{\text{interior}} \leftarrow S^{\text{interior}} \cup \{ & \mathbf{b}_i, \mathbf{b}_{i+1}, (\mathbf{b}_i(1), \mathbf{b}_i(1)+1), \\
& \ldots, (\mathbf{b}_{i+1}(1)-1, \mathbf{b}_{i+1}(1)), (\mathbf{b}_{i+1}(2), \mathbf{b}_{i+1}(2)+1), \\
& \ldots, (\mathbf{b}_{i+1}(1)-1, \mathbf{b}_{i+1}(1))\},
\end{aligned}
$$

where all edges are phosphodiester bonds, except the hydrogen bonds $\mathbf{b}_i, \mathbf{b}_{i+1}$.

- Bifurcation loop: Let $\mathbf{z}$ represent the number of base pairs in the loop. If there are more than two base pairs (i.e. $\mathbf{z} > 2$) in this loop, then

$$
\begin{aligned}
S^{\text{bifurcation}} \leftarrow S^{\text{bifurcation}} \cup \{ & \mathbf{b}_i, \mathbf{b}_{i+1}, \ldots, \mathbf{b}_{i+\mathbf{z}-1}, \\
& (\mathbf{b}_i(2), \mathbf{b}_i(2)+1), \ldots, (\mathbf{b}_{i+1}(1)-1, \mathbf{b}_{i+1}(1)), \\
& (\mathbf{b}_{i+1}(2), \mathbf{b}_{i+1}(2)+1), \ldots, (\mathbf{b}_{i+2}(1)-1, \mathbf{b}_{i+2}(1)), \\
& \ldots, \\
& (\mathbf{b}_{i+\mathbf{z}-2}(2), \mathbf{b}_{i+\mathbf{z}-2}(2)+1), \ldots, (\mathbf{b}_{i+\mathbf{z}-1}(1)-1, \mathbf{b}_{i+\mathbf{z}-1}(1)) \\
& (\mathbf{b}_{i+\mathbf{z}-1}(2), \mathbf{b}_{i+\mathbf{z}-1}(2)+1), \ldots, (\mathbf{b}_i(1)-1, \mathbf{b}_i(1))\},
\end{aligned}
$$

where all edges are phosphodiester bonds, except the hydrogen bonds $\mathbf{b}_i, \mathbf{b}_{i+1}, \ldots, \mathbf{b}_{i+\mathbf{z}-1}$.

- **Step 2 Obtain the sequence of each motif:** The sequence of the motif includes all the paired nucleotides and unpaired nucleotides in the motif that are aligned in order of index.

#### Details of Nucleotide Localized Information Matrix

After utilizing the one hot encoding to represent the categorical variables, the size of the nucleotide localized information matrix becomes $L \times 18$, where 4 columns are for 4-elements nucleotide one hot encoding, 12 columns are for two 6-elements motif one hot encoding, and 2 columns are for free energy of two motifs.

#### Utilizing Cumulative Distribution Function of Normal Distribution to Rescale the Free Energy of Motifs

Different from the binary valued features from the categorical variables, the free energy of motifs are numerical in nature. To prevent learning difficulties, we rescale these values to the $[0, 1]$ interval by utilizing the cumulative distribution function (CDF) of normal distribution with $\mu$ and $\sigma$, where $\mu$ is the mean of normal distribution and $\sigma$ is the standard deviation of normal distribution. Here, the reason why we utilize CDF of normal distribution to rescale the free energy is that the rate of CDF of normal distribution converging to 0 and 1 can be controlled by the parameter $\sigma$. We wish the scaling function converging to 0 and 1 neither too fast nor too slow. In other words, we expect to customize a scaling function so that it can be more sensitive to the different free energy values. The nature of CDF of normal distribution exactly satisfies this requirement.

[2] Sergey Ioffe and Christian Szegedy. Batch normalization: Accelerating deep network training by reducing internal covariate shift. *arXiv*, 2015.

[3] Marcin Magnus, Maciej Antczak, Tomasz Zok, Jakub Wiedemann, Piotr Lukasiak, Yang Cao, Janusz M Bujnicki, Eric Westhof, Marta Szachniuk, and Zhichao Miao. Rna-puzzles toolkit: a computational resource of rna 3d structure benchmark datasets, structure manipulation, and evaluation tools. *Nucleic acids research*, 48(2):576–588, 2019.

[4] Vinod Nair and Geoffrey E Hinton. Rectified linear units improve restricted boltzmann machines. *ICML*, pages 807–814, 2010.
